# The BXD Mouse Strains are a Model System for Studying Optic Nerve Regeneration

**DOI:** 10.1101/204842

**Authors:** Jiaxing Wang, Ying Li, Rebecca King, Felix L. Struebing, Eldon E. Geisert

## Abstract

The present study is designed to identify the influences of genetic background to optic nerve regeneration using the two parental strains C57BL/6J and DBA/2J and 7 BXD recombinant inbred strains. To study regeneration in the optic nerve, *Pten* was knocked down in the retinal ganglion cells using AAV, and a mild inflammatory response was induced by an intravitreal injection of zymosan with CPT-cAMP, and the axons were damaged by optic nerve crush (ONC). Regenerating axons were labeled by Cholera Toxin B and quantified 14 days after ONC. The number of axons at 0.5 mm and 1 mm from the crush site were counted. In addition, we measured the distance that 5 axons had grown down the nerve and the longest distance a single axon reached. Results showed a considerable amount of differential axonal growth across all 9 BXD strains. There was a significant difference (*P*=0.014 Mann-Whitney U test) in the regenerative capacity in the number of axons reaching 0.5 mm from a low of 1487.6 ± 264.9 axons in BXD102 to a high of 4175.8 ± 648.6 axons in BXD29. There were also significant differences (*P*=0.014 Mann-Whitney U test) in the distance axons traveled, looking at a minimum of 5 axons with the shortest distance was 787.2 ± 46.5µm in BXD102 to a maximum distance of 2025.5 ± 223.3µm in BXD29. These results reveal that genetic background can modulate axonal regeneration and that the BXD strains are a particularly well-suited model system.

## INTRODUCTION

Over the last decade, significant advances have been made in approaches to induce regeneration of retinal ganglion cell (RGC) axons through the optic nerve[1–5]. The regeneration and survival of RGCs is influenced by interactions between multiple cellular processes (for review see [5–7]). The number of genes and molecular pathways that modulate the regenerative response in the mammalian optic nerve reveals that induced axonal regeneration (or the lack of regeneration in the normal adult CNS) is a complex trait[1, 2, 8–11]. Complex traits are controlled by multiple genomic elements, some of which are associated with specific molecular functions and others are believed to be associated with more generalized cellular functions[12–14]. This complexity of axonal regeneration could be predicted for we know that successful regeneration involves multiple cellular processes. The first is the survival on the injured retinal ganglion cell involving modulating apoptosis[15, 16], autophagy[1] and response to growth factors[11, 17-19]. The second necessity for axonal regeneration to occur is the growth of the axon itself down the optic nerve. This includes distinct pathways associated with the axon growth program[20]. The third series of events may be directly related to cellular elements that inhibit axonal growth in the adult CNS that are glial in origin, involving astrocytes[21, 22], oligodendrocytes[10] or the glial scar[21, 22]. Our goal is to take a systems biology approach to the study of optic nerve regeneration and to treat the process as a complex trait.

Our working hypothesis is that current regeneration treatments can be influenced by genetic background and within that genetic background are specific genomic elements that can be identified. Our group has used a systems biology approach working with the BXD recombinant inbred strains of mice to define genomic elements affecting the response of the retina to optic nerve damage[23] and to blast injury[24]. The power of the BXD strain set derives from the shuffled genomes of the parental strains (C57BL/6J mice and the DBA/2J mice). Both of the parental strains are fully sequenced and there are over 4.8 million known single nucleotide polymorphisms, deletions, and insertions between them. In the first 102 BXD strains, there are over 7000 break-points in the genomes between the parental strains. All of the BXD strains are fully mapped. This allows for a rapid mapping of phenotypic data onto genomic elements to define loci modulating the phenotype in a quantitative trait analysis [25, 26]. All of these information and powerful bioinformatic tools are available on the GeneNetwork website (<u>www.genenetwork.org</u>) and are used to define the complex genetics underlying induced regeneration in the optic nerve.

We use the BXD recombinant inbred strains to examine the regeneration response 14 days after optic nerve crush in mice that have received a previously described[1, 3] regeneration treatment of knocking down *Pten* (phosphatase and tensin homolog) and intravitreal injection of zymosan and CPT-cAMP. Regenerative response is determined by defining the number of axons regenerating as well as the distance these axons have traveled.

## MATERIALS AND METHODS

### Mice

Nine strains including seven BXD recombinant inbred strains and their parental strains – C57BL/6J and DBA/2J were used in this study. All of the mice were 60-70 days of age at the time of initial treatment (See supplemental Table 1). The mice were housed in a pathogen-free facility at Emory University, maintained on a 12-hour light/dark cycle, and provided with food and water ad libitum. All procedures involving animals were approved by the Animal Care and Use Committee of Emory University and were in accordance with the ARVO Statement for the Use of Animals in Ophthalmic and Vision Research. Controls were run with the C57BL/6J (n=6) and DBA/2J (n= 6) mice strains. For the studies of regeneration, we examined axon growth in the parental strains, C57BL/6J (n=5) and DBA/2J (n=8); along with, seven BXD strains: BXD11 (n=5), BXD29(n=4), BXD31(n=4), BXD38(n=4), BXD40(n=9), BXD75(n=5) and BXD102(n=5).

**Table 1.** Genetic characteristics of genes that are known to affect optic nerve regeneration between C57BL/6J and DBA/2J mice.

|  |  |
| --- | --- |
| **Gene with cis-QTL** | *Fgf2*[39] |
| **Genes with Non-synonymous SNPs** | *Mapk10* [40], *Rtn4*[41, 42], *Ctgf*[11], *Tlr2*[43], *Rock1*[44],<br>*Rock2*[44, 45], *Clec7a*[43], *Csf2*[46] |
| **Other investigated genes**<br>(Genes that are found to have neither cis-QTL nor Non-synonymous SNPs ) | *Braf*[47, 48], *Bcl2*[9], *Myc*[49], *Cntf*[50], *Dpysl2*[51],<br>*Ecel1*[52], *Map3k12*[53], *Egfr* [17], *Gsk3b*[54, 55], *Hhex*<br>[56], *Hnrnpk* [57], *Spp1*[58], *Stat3*[2, 59, 60], *Klf4*[2, 60,<br>61], *Klf6*[2, 61, 62], *Klf7*[2, 61, 62], *Klf9*[2, 61], *Lif*[50, 63],<br>*Rtn4r*[10, 64, 65], *Ntn1*[66, 67], *Pten*[1], *Ptprs*[68],<br>*Rhoa*[69-71], *Cxcl12*[72], *Set*[73], *Socs3*[37, 74], *Sox11*[5,<br>75], *Tet1*[76], *Tet3*[76], *Wnt10b*[77], *Slc30a3*[78], *Bag1*<br>[79] |

### Surgery

The optic nerve regeneration protocol developed by others[1, 3, 4] was used to induce regeneration after optic nerve crush (ONC). The treatment included knocking down of *Pten* using AAV-shPTEN-GFP, and intravitreal injection of Zymosan plus CPT-cAMP. We followed a similar protocol with minor modifications. One is that we used AAV-shPTEN-GFP (*Pten* short hairpin RNA-GFP packaged into AAV2 backbone constructs, titer = 1.5×10^12^ vg/ml) to knock down *Pten* instead of cre recombinase-mediated knock-out in *Pten*-floxed mice. The shRNA target sequence is 5’-AGGTGAAGATATATTCCTCCAA-3’ as described by Zukor K et al[27]. Our immunostaining also proved an efficient suppression of *Pten* expression in the retina ganglion cells by this *Pten* shRNA (Supplementary Figure 1). Two weeks prior to ONC, the mice were deeply anesthetized with 15 mg/kg of xylazine and 100 mg/kg of ketamine and injected intravitreally with 2µL of AAV-shPTEN-GFP. Optic nerve crush was performed as described by Templeton and Geisert[28]. Briefly, the mice were deeply anesthetized with a mixture of 15 mg/kg of xylazine and 100 mg/kg of ketamine. Under the binocular operating scope, a small incision was made in the conjunctiva, the optic nerve was visualized and then crushed 1 mm behind the eye with angled crossover tweezers (Dumont N7) for 5 seconds, avoiding injury to the ophthalmic artery. Immediately following ONC, Zymosan (Sigma, Z4250, Lot#BCBQ8437V) along with the cAMP analog CPT-cAMP (Sigma, C3912, Lot#SLBH5204V) (total volume 2µL) were injected into the vitreous to induce an inflammatory response and augment regeneration. The animals were allowed to recover from anesthesia and returned to their cages. Twelve days after ONC (2 days before sacrifice) the animals were deeply anesthetized and Alexa Fluor^®^ 647-conjugated Cholera Toxin B (CTB) (ThermoFisher, C34778) was injected into the vitreous for retrograde labeling of the regenerated axons. All the intravitreal injections and optic nerve crushes were done by one well-trained postdoctoral fellow to avoid technical variation during the surgical procedure. At 14 days after ONC the mice were deeply anesthetized and perfused through the heart with phosphate buffered saline (pH 7.3) followed by 4% paraformaldehyde in phosphate buffer (pH 7.3).

### Preparation of the optic nerve

Optic nerves along with optic chiasm, and brains were dissected and post fixed in 4% paraformaldehyde in phosphate buffer overnight. The optic nerve was cleared with FocusClear™ (CelExplorer, Hsinchu, Taiwan) for up to 4 hours until totally transparent. A small chamber was built on the slide to provide enough space for the whole nerve thickness and to keep the nerve from being damaged from flattening. The optic nerve was then mounted in the chamber using MountClear™ (CelExplorer, Hsinchu, Taiwan) and the slides were cover-slipped.

FocusClear has been used to clear brain tissue for whole brain imaging [29] as well as clearing of the optic nerve of transgenic zebrafish to observe axon regeneration [30]. It allows us to scan the whole thickness of the optic nerve for better understanding of the status of axon regeneration. It provides clear imaging of regenerated axons from the optical slices scanned by confocal microscope for counting. It also allowed us to determine the longest 5 axons or single axon growth along the nerve from z-stack of the whole nerve (Figure 1).

**Figure 1.**
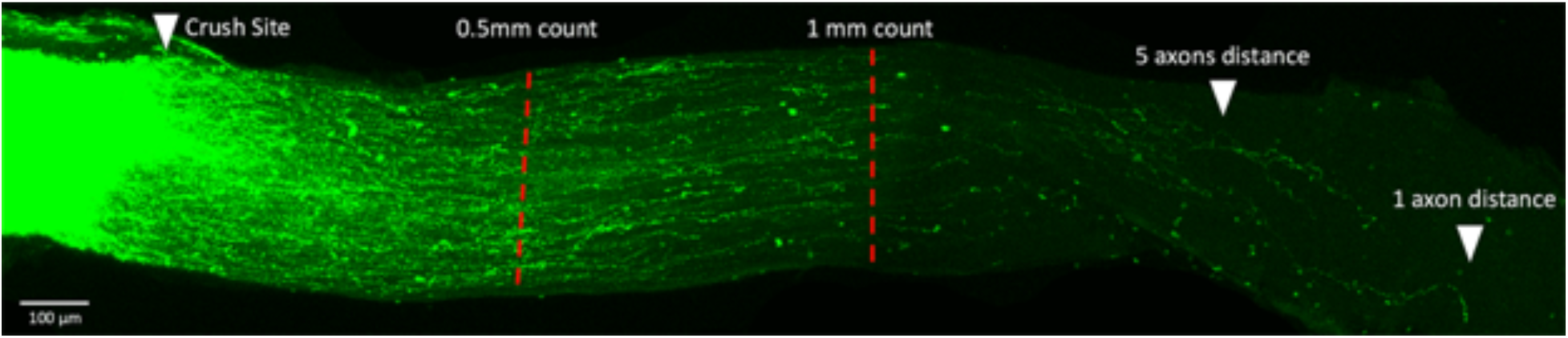
The regenerating axons in the optic nerve 14 days after optic nerve crush. We have indicated the regions of the nerve where axons were counted as well as the distance that 5 axons or 1 axon regenerated down the nerve. Scale bar represents 100 µm.

### Quantitation of axon regeneration

Cleared optic nerves were examined on a confocal microscope by scanning through individual optical slices. Green pseudo-color was used for CTB-labeled axons in all the optic nerve images of this study for clear visual observation. Stacked images were taken at 5 µm increments, a total of 50-100 optical slices for each optic nerve. The number of CTB labeled axons at 0.5 mm from the crush site were counted in at least 6 sections per case and calculated by the equation (Σad= πr^2^ ∗ [average axons/mm]/t) as described by Leon et al. in 2000 [31]. The cross-sectional width of the nerve was measured at the point at which the counts were taken and was used to calculate the number of axons per millimeter of nerve width. The total number of axons extending distance *d* in a nerve having a radius of *r*, was estimated by summing over all sections. Since we used confocal images instead of longitudinal cross sections described in previous studies[3, 4], the optical resolution in *z* (0.5µm) was considered as the *t* (thickness of the slide) in the equation. In some strains very few axons were observed 1mm from the crush site, and for this reason we used direct counts of axons as a measure of regeneration. We also measured the distance that 5 axons had grown down the nerve and the longest distance a single axon reached for each nerve from z-stack image of the whole nerve.

### Transfection Efficiency of AAV-shPTEN-GFP

C57BL/6J mice (N=4) and BXD29 mice (N=4) were deeply anesthetized with a mixture of 15 mg/kg of xylazine and 100 mg/kg of ketamine and injected with 2 µl of AAV-shPTEN-GFP into the left eye. Two weeks later, they were deeply anesthetized as described above and perfused through the heart with saline followed by 4% paraformaldehyde in phosphate buffer (pH 7.3). For the retinal flat mounts, the retinas were removed from the globe and rinsed in PBS with 1% Triton X-100, blocked in 5% Bovine Serum Albumin (BSA) 1-hour room temperature and placed in primary antibodies RBPMS (Millipore, Cat. # ABN1376) at 1:1000 and GFP (Novus Biologicals, CAT #NB100-1770) at 1:1000 4°C overnight. The retinas were rinsed with PBS and placed in secondary antibodies, (Anti-Goat IgG(H+L) CFTM 488A, Sigma, Cat. #SAB4600032 and Alexa Fluor 594 AffiniPure Donkey Anti-Guinea, Jackson Immunoresearch, Cat. #706-585-1148) at 1:1000 for 1 hour at room temperature. After three washes of 15 minutes each, retinas were flat mounted and cover-slipped using Fuoromount - G (Southern Biotech, Cat. #0100-01) as a mounting medium. Four confocal images were taken in each quadrant at 2 mm away from the optic nerve of each retina. Four retinas from 4 mice of each strain were included. Cell number were determined manually by using the cell counter in ImageJ. RBPMS was used as a marker to label the total number of RGCs [32, 33]. Transfection Efficiency are calculated as the number of AAV transfected RGCs (GFP^+^RBPMS^+^ cells) divided by total number of RGCs (RBPMS^+^ cells).

### Bioinformatic analysis of known regeneration genes in the BXD strain set

By searching the literature, we generated a list of genes that are known to have effects on optic nerve regeneration either directly or indirectly (Table 1). All those genes were examined for high Likelihood Ratio Statistics (LRS) scores and cis Quantitative Trait Loci (cis-QTLs) using the GeneNetwork database. They were then put into the SNP (Single Nucleotide Polymorphism) browser of GeneNetwork as well as UCSC genome browser (Mouse, GRCm38/mm10) to identify non-synonymous SNPs between C57BL/6J and DBA/2J. All the identified rsIDs of non-synonymous SNPs were then put into Ensembl (www.ensembl.org/) for SIFT analysis [34] to predict whether the SNP affects protein function.

### Statistical Analysis

Data are presented as Mean ± SE (Standard Error of the Mean). Differences in axon counts and regeneration distance were analyzed by Mann-Whitney U test. The differences in transfection efficiency were analyzed by 2-independent sample t-test using SPSS (Statistical Package for the Social Sciences) statistical package 24.0 (SPSS, IBM, Chicago, IL, USA). A value of *p*<0.05 was considered statistically significant.

## RESULTS

### Genetic Background Modulates RGC Axon Regeneration

In the present study, we examined the effects of genetic background on the regenerative response of retinal ganglion cells. The BXD recombinant inbred strains were chosen as a genetic reference panel due to distinct advantages these strains have to offer. The first advantage is there are over 80 well characterized strains of mice available through Jackson laboratories. The current set of strains is far larger than any other mature RI resource and will now allow for sub-megabase mapping resolution. The second benefit of the BXD strains are the large microarray datasets specifically for the eye (HEIMED database[14]; HEI Retina Database [35]; Optic Nerve Crush Database[23]; and the DoD Normal Retina Database[36]) along with the numerous ocular phenotypes RGC numbers, IOP, eye size, retinal area, etc[24]. GeneNetwork also offers an array of highly interactive series of bioinformatic tools that aid in the analysis of data generated with the BXD strains.

Regeneration of axons in the optic nerve was examined in nine strains of mice: the two parental strains (C57BL/6J and DBA/2J mice) and seven BXD strains (BXD11, BXD29, BXD31, BXD38, BXD40, BXD75 and BXD102) (Figure 2). As an internal control, we examined the ability of untreated retinas to regenerate following optic nerve crush in both the C57BL/6J mouse and the DBA/2J mouse (Figure 2). For all mice, the axonal regeneration was evaluated 14 days following optic nerve injury. In the two strains of the control group (C57BL/6J and DBA/2J) there was no detectable axonal regeneration; while in all of the strains receiving the regeneration treatment there was a significant regenerative response, both in the number of axons counted at 0.5mm and 1mm (Figure 4), as well in the distance the axons traveled (Figure 5). These data demonstrate that the regeneration treatment, knocking down *Pten* and inducing a mild inflammation by injecting of zymosan and CPT-cAMP, produces significantly more regenerating axons than observed in control animals that did not receive treatment.

**Figure 2.**
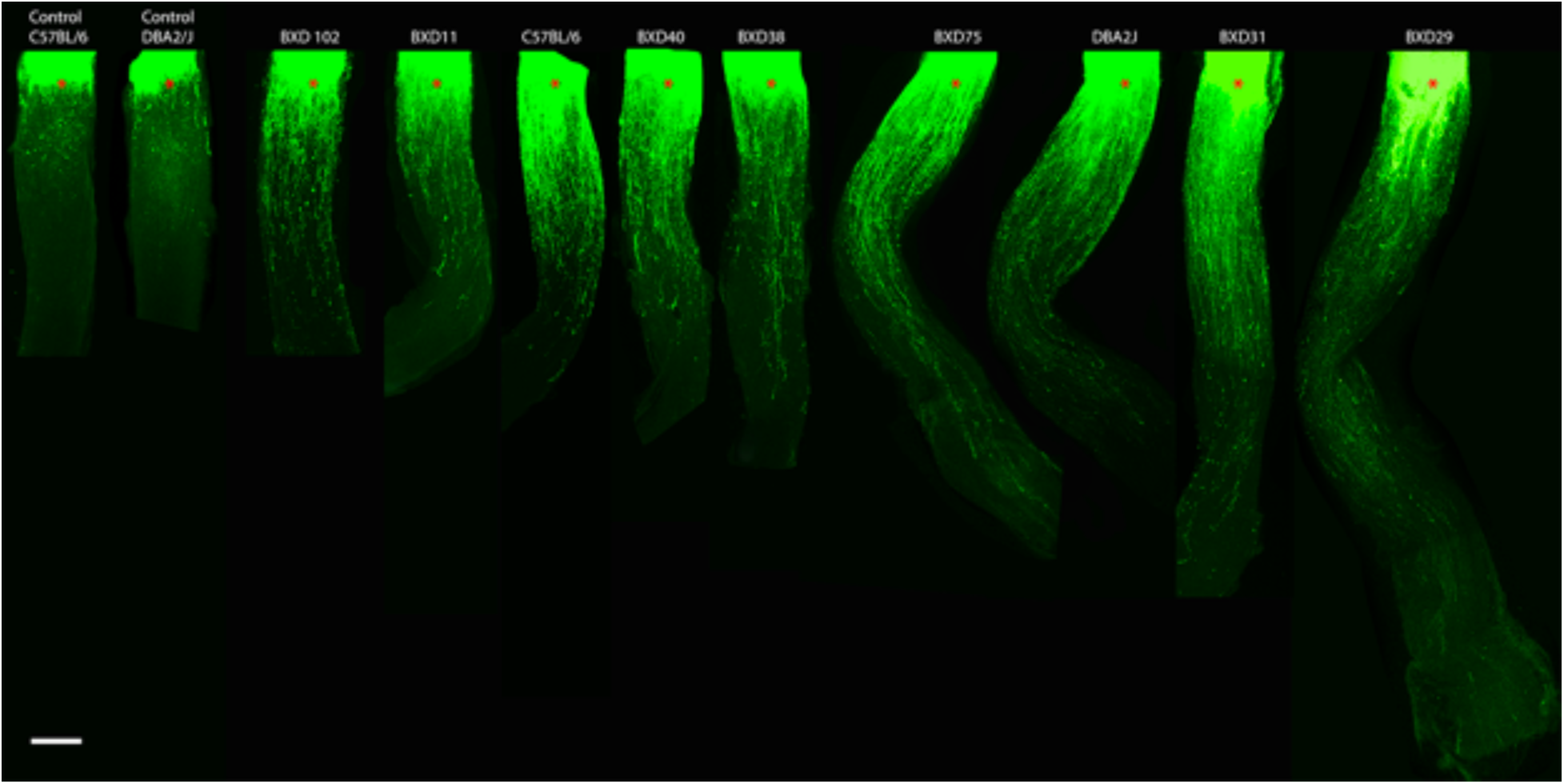
Genetic background affects regenerating axons in the optic nerve following crush. The figure is a series of photomicrographs from 11 optic nerves selected from 9 different strains of mice. The first two images on the far left are from control mice that did not receive the regeneration treatment prior to optic nerve crush (Control C57BL/6J and Control DBA/2J). All of the remaining nerves were from animals in which *Pten* was knocked-down and a mild inflammatory response was induced. The strain with the least regeneration was BXD102 and the strain with the greatest regeneration was BXD29. Red asterisks represent the crush site. The scale bar represents 200µm.

**Figure 3.**
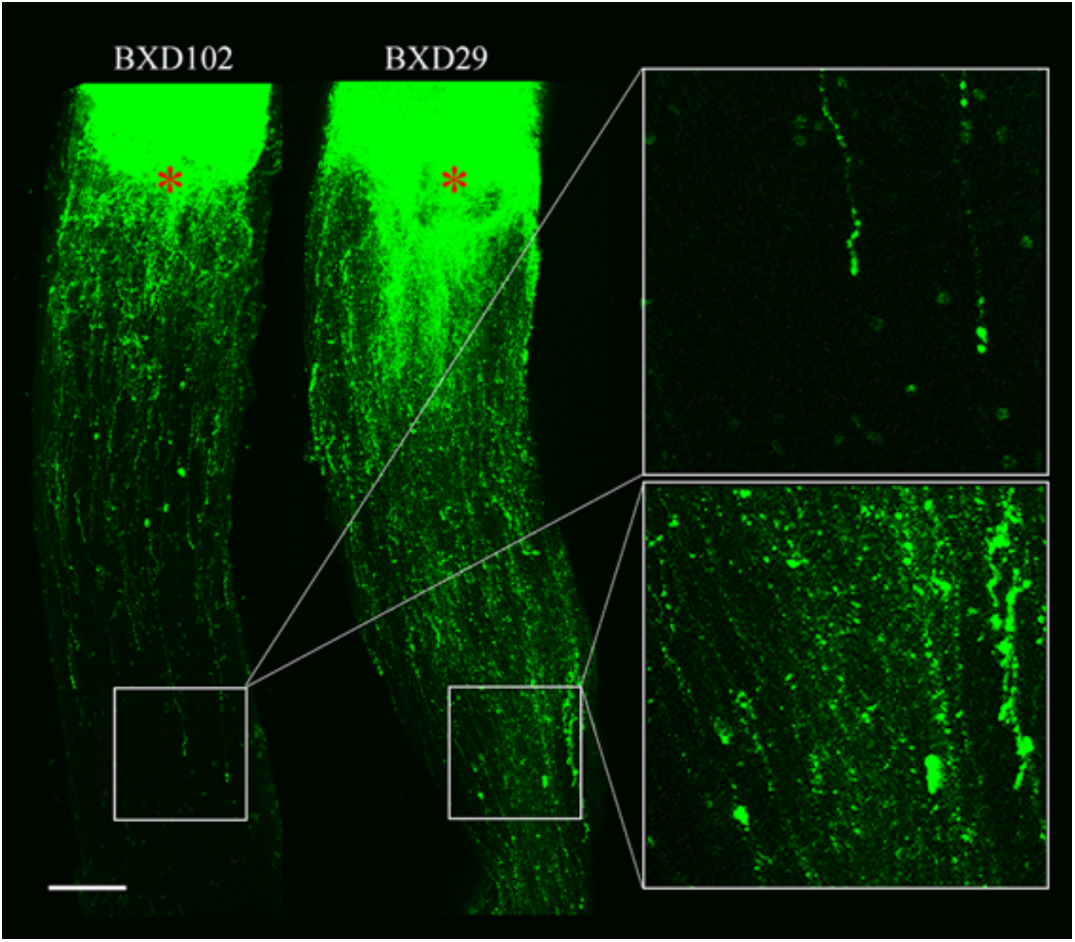
Comparison of regenerated axons in strains with the least regeneration (BXD102) and the greatest regeneration (BXD29). Higher magnification of axons at 1mm (Boxed region) from crush site were shown relatively. The scale bar represents 100µm.

**Figure 4:**
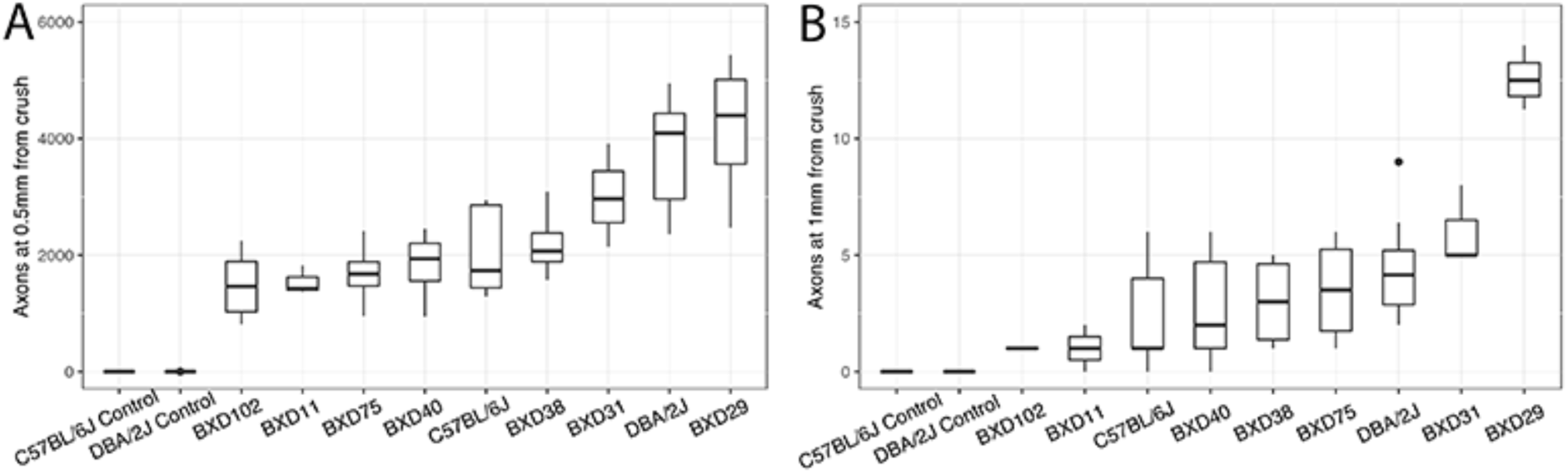
The number of axons at 0.5mm (A) and 1mm (B) from the crush site in two control strains (DBA/2J and C57BL/6J untreated mice) and in 9 strains treated with the regeneration protocol. Boxplots show median, 25^th^ and 75^th^ percentile, maximum, and minimum values for each BXD RI strain.

**Figure 5:**
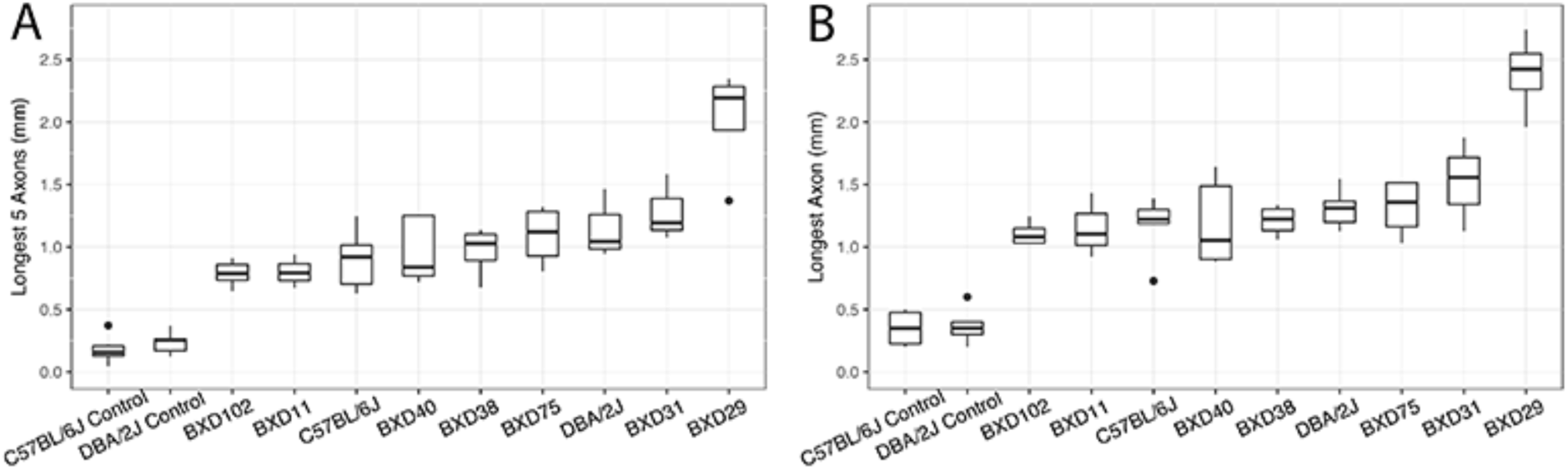
The longest distance that 5 axons regenerated (A) and the longest regeneration for a single axon (B) are shown for the two control strains (DBA/2J and C57BL/6J untreated mice) and in 9 strains treated with the regeneration protocol. Boxplots show median, 25^th^ and 75^th^ percentile, maximum, and minimum values for each BXD RI strain.

The number of axons at 0.5mm and 1mm distal to the crush, is an estimate of the influence of genetic background on the total regenerative effect of the treatment. As can be seen in Figure 2 and Figure 4, there is a considerable difference between strains on the total number of axons reaching 0.5mm and 1.0mm. The strain with the least number of axons in both cases was BXD 102. At both distances from the crush site, the strain with the greatest number of axons was BXD29. The difference was significant (*P*=0.014 Mann-Whitney U test) in the number of regenerated axons reaching 0.5 mm (from a low of 1487.6 ± 264.9 axons in BXD102 to a high of 4175.8 ± 648.6 axons in BXD29) from the crush site. There was also a significant difference (*P*=0.007 Mann-Whitney U test) in the number of axons at 1mm, from a low of 1±0 axons in BXD102 to a high of 12.6 ± 0.6 axons in BXD29. The two strains that displayed the least and most robust (BXD102 and BXD29) regenerative response are illustrated in Figure 3.

The total length of a regenerating axon was also measured. This measure may provide an estimate of the rate at which the axon can grow down the injured optic nerve. When we examined axon length there was also a clear difference in growth across the BXD strains (Figure 2, Figure 5). In the control animals, virtually no regenerating axons were observed. When we examined the distance a minimum of 5 axons traveled, a significant difference was observed across the BXD strains. The strain with the shortest regenerating 5 axons was BXD102 with a mean distance of 787.2 ± 46.5µm, and the strain with the longest group of 5 axons was BXD29 with a mean distance of 2025.5 ± 223.3µm (*P*=0.014 Mann-Whitney U test). A similar result was observed when examining the distance of the longest single axon traveled in the nerve, with BXD102 having the shortest average distance (1107 ± 40.6µm) and BXD29 the longest (2386.8 ± 162.6µm, *P*=0.014 Mann-Whitney U test). Thus, the ability of axonal regeneration (both the number of regenerating axons and the distance traveled) is affected by the genetic backgrounds in the BXD strains with BXD102 having the least regeneration and BXD29 having the most robust regenerative response. These data revealed that genetic background can have a striking effect on the regenerative capacity of axons within the optic nerve.

### Transfection Efficiency of AAV-shPTEN-GFP

One possible explanation for the difference is axonal regeneration is a differential transfection of the retinas from strain to strain by the AAV-shPTEN-GFP vector. To control for this possibility, we examined the transfection efficiency and level of *Pten* knock down in the strain with the most robust axon regeneration (BXD29) and another strain with moderate axon regeneration (C57BL/6J). There was no statistically significant difference between the two strains. For the BXD29 strain (n=4) the mean transfection rates was 51.6%±1.3% and for the C57BL/6J strain (n=4) the mean transfection rate was 53.3% ± 1.7% (Figure 6), indicating that the difference of regeneration response is not due to different transfection efficiency.

**Figure 6.**
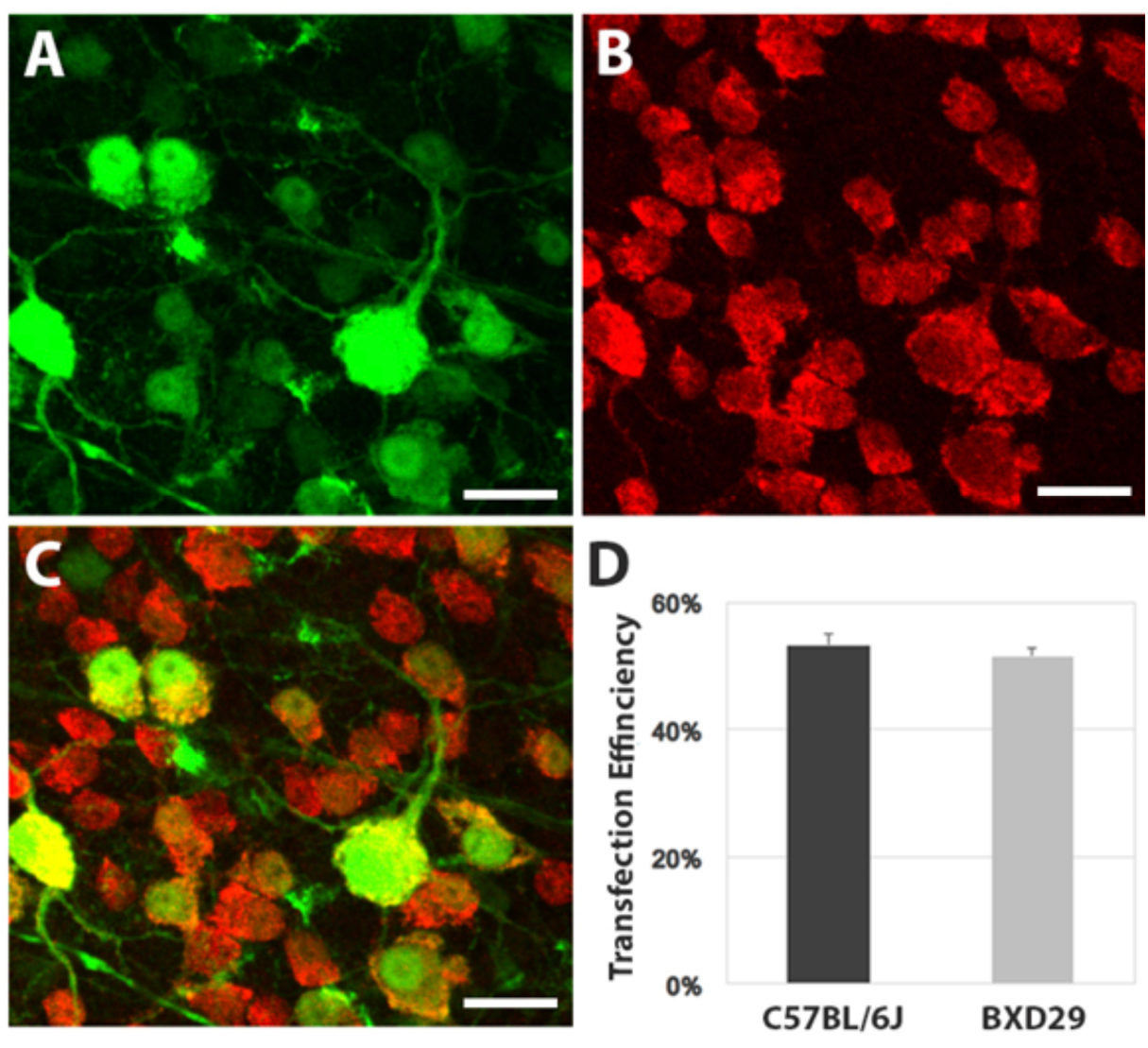
Transfection Efficiency of AAV-shPTEN-GFP. AAV transfected cells are labeled by GFP in green (A) and total number of RGCs are labeled by RBPMS in red (B). Merged channel shown in (C). No statistical difference of transfection efficiency was found between C57BL/6J mice and BXD29 mice (D). Scale bar represents 100 µm.

### Potential Contribution of Known Genes Affecting Axon Regeneration BXD Strains

Previous studies have identified a number of genes that affect the ability of axons to regenerate in the injured optic nerve (Table 1). Using the bioinformatic tools on GeneNetwork, it is possible to define any of the regeneration associated genes that are either differentially expressed forming a cis-QTL in the BXD strains or having non-synonymous SNPs between C57BL/6J and DBA/2J mice.

The cis-QTL is a QTL that maps to the location of the gene that produces the mRNA or protein. We usually use the LRS (Likelihood Ratio Statistic) score to represent the association or linkage between differences in traits and differences in particular genotype markers or specific genes. While a statistically significant cutoff can only be determined through permutation tests, LRS scores of >17 usually approximate the significance threshold of *p*<0.05 and are worthy of attention[14]. If a cis-QTL has a high LRS score, it is considered that this genetic locus is strongly linked to a certain phenotype and is able to influence the phenotype by regulating this locus. In other words, change of the expression level of this gene will have a higher chance to alter the phenotype, which in our case, is the axonal regeneration. In this process, two regeneration associated genes were identified with cis-QTLs, *Fgf2* and *Klf9* (Table 1). Only one of these cis-QTLs, *Fgf2*, is valid. The other, *Klf9,* contained a difference in the genetic sequences between C57BL/6J and DBA/2J mice at the exact site where the microarray probe binds. This difference in sequence will lead to differential binding of the probe and a false positive LRS score. Thus, there was one cis-QTL (*Fgf2*) present in the BXD strains that could potentially affect the regenerative response.

We also examined the BXD strains to define genes with non-synonymous SNPs. A non-synonymous SNP between the parent strains (C57BL/6J and DBA/2J) is potentially able to alter the protein structure and function, ultimately leading to the different phenotype. The BXD strains that inherited different alleles may also have different phenotypes. There were eight genes (*Mapk10, Rtn4, Ctgf, Tlr2, Rock1, Rock2, Clec7a and Csf2)* with non-synonymous SNPs between C57BL/6J and DBA/2J. The SIFT analysis[34] revealed that only two of the eight genes, *Mapk10* (JNK3) and *Rtn4* (NOGO), had SNPs that was predicted to likely affect protein structure/function (rs36844177 in *Mapk10,* SIFT=0.01 and rs29465940 in *Rtn4*, SIFT=0.03). Thus, in the BXD strain set, only three genes known to be associated with axonal regeneration, *Fgf2, Mapk10* (JNK3), and *Rtn4* (NOGO), are actively different between C57BL/6J and DBA/2J mice and potentially contribute to the different response of axonal regeneration across the BXD strains.

## DISCUSSION

Over the past several years, advances in optic nerve axon regeneration have taken what was once thought of as an unachievable goal to the point where axonal regrowth after injury is a reality. Several different protocols are being used to promote axonal regeneration[2, 5, 37]. In the present study, we chose a popular protocol developed by others[3, 4] that involves knocking down *Pten* and causing a mild inflammatory response. The BXD recombinant strains are ideal for testing the effects of genetic background with the protocol of knocking down *Pten*, for there are not significant differences in *Pten* between the C57BL/6J strain and the DBA/2J strains. There are no non-synonymous SNPs found in the *Pten* gene between C57BL/6J and DBA/2J mice. Furthermore, there is a similar level of expression of *Pten* mRNA across all of the BXD strains in the DoD normal retina datasets housed on GeneNetwork. We also examined transfection efficiency in two strains (BXD29 and C57BL/6J) that respond differently to the regeneration treatment. There was no difference in transfection efficiency between these two strains. Thus, the difference in the axon regeneration we observed between the different BXD strains cannot be explained by the expression levels of *Pten* in the strains or a differential level of transfection by the AAV2 vector. This leaves only one possibility that the difference in axonal regeneration we observed is due to the specific segregation of genomic elements cross the BXD strains.

Using the BXD strains we were able to demonstrate the effect of genetic background on the regenerative capacity of axons in the optic nerve. In all strains tested, the amount of regeneration was considerably greater than that observed in mice that did not receive the *Pten*/Zymosan/cAMP treatment. The regeneration responses of C57BL/6J mice we observed were not as strong as described in other studies, a possible reason could be that we were using AAV-shPTEN-GFP to knock down *Pten* instead of cre recombinase-mediated *Pten* knock out. The other factor to be noticed is that the age of mice we used are over 60 days at the time of initial treatment, much older than reported in other studies[3, 4]. This also provides strong evidence that the regeneration response can happen not only in young adult mice but also in older mice. Among the strains treated to promote regeneration, some strains, like BXD102, showed a modest regenerative response; while other strains, like BXD 29, consistently demonstrated a high number of regenerating axons and axons that traveled longer distances down the injured optic nerve. When the response of the parental strains C57BL/6J and DBA/2J was compared to the BXD strains with extreme regenerative responses, there was a clear indication of genetic transgression. If we look at the number of axons at 0.5mm and 1mm from the crush site (Figure 4), there are BXD strains that have fewer axons and BXD strains that have more axons than the parental strains. This difference in regeneration is indicative of genetic transgression. These data reveal that it is not a single genomic locus causing the variability in the regenerative response, for if that were the case the extremes would be similar to the parental strains. This is a clear indication that multiple genomic loci are segregating across the BXD strains to affect the regenerative response of the optic nerve axons. Thus, axon regeneration is a complex trait with multiple modulating genomic loci in the BXD strains.

A complex trait could be driven by a handful of protein coding genes as well as noncoding variants that presumably affect gene regulation[38]. Multiple genes have been identified in recent years to have an impact on axon regeneration (Table 1). It is possible that some or all of these pathways are varying in the BXD strains and are influencing the outcome of the induced regeneration observed in the present study. With the DoD normal retina datasets[36] and the bioinformatic tools hosted on GeneNetwork, we examined all of the known regeneration-related genes to determine if they were able to modulate the regenerative response across the strains by having either cis-QTLs (differentially expressed genes) or non-synonymous SNPs that would affect protein function. Of all of the known genes known to alter the regenerative response of optic nerve axons, only two were potential candidates for modulating regeneration in the BXD strains. The only cis-QTL in the list of genes was *Fgf2* (LRS = 67.8). SNP analysis identified 8 genes with nonsynonymous SNPs in the list of regeneration genes (Table 1). All of the nonsynonymous SNPs were examined using a SIFT analysis and only 2 SNPs (rs36844177 in *Mapk10*(JNK3) and rs29465940 in *Rtn4*(NOGO)) was predicted to alter protein function. Therefore, three possible genomic elements that could be affecting regeneration in the BXD strains are *Fgf2* (Chr3, 37.3 Mb), *Mapk10* (Chr5, 102.9Mb) and *Rtn4* (Chr11, 29.7Mb). Beyond all these, we believe that there are still unknown genomic elements modulating the regenerative response of the axons.

In conclusion, the ability of optic nerve regeneration response to injury is different across BXD strains given different genetic background. Quantitative trait analysis may provide us with new insights to axon regeneration, and maybe new loci of novel genes or non-coding elements that are involved in axon regeneration. The BXD strains are a good model system and a useful genetic reference panel to study the regeneration of axons in the optic nerve.

## Acknowledgements

This study was supported by an Unrestricted Grand from Research to Prevent Blindness, NEI grant R01EY017841 (E.E.G.), Owens Family Glaucoma Research Fund, P30EY06360 (Emory Vision Core) and DoD CDMRP Grant W81XWH-12-1-0255 from the USA Army Medical Research & Materiel Command and the Telemedicine and Advanced Technology (E.E.G.).

## Supplementary data

### Supplementary Method

### Evaluation of *Pten* knock down by immunofluorescence staining

Six C57BL/6J mice were deeply anesthetized with a mixture of 15 mg/kg of xylazine and 100 mg/kg of ketamine and perfused through the heart with saline followed by 4% paraformaldehyde in phosphate buffer (pH 7.3). For the retinal flat mounts, the retinas were removed from the globe and rinsed in PBS with 1% Triton X-100, blocked in 5% Bovine Serum Albumin (BSA) 1 hour room temperature and placed in primary antibodies *Pten* (Cell Signaling, Cat. # 9559) at 1:200 and GFP (Novus Biologicals, CAT #NB100-1770) at 1:1000 4°C overnight. The retinas were rinsed with PBS and placed in secondary antibodies, (Anti-Goat IgG(H+L) CFTM 488A, Sigma, Cat. #SAB4600032 and Alexa Fluor 594 AffiniPure Donkey Anti-Rabbit, Jackson Immunoresearch, Cat. #711-585-152) at 1:1000 and To-PRO-3 Iodide (Molecular Probes, Cat. # T3605) was applied 1:1000 as a nuclear counterstain for 1 hour at room temperature. After three washes of 15 minutes each, retinas were flat mounted and cover-slipped using Fuoromount - G (Southern Biotech, Cat. #0100-01) as a mounting medium.

**Supplementary figure 1.**
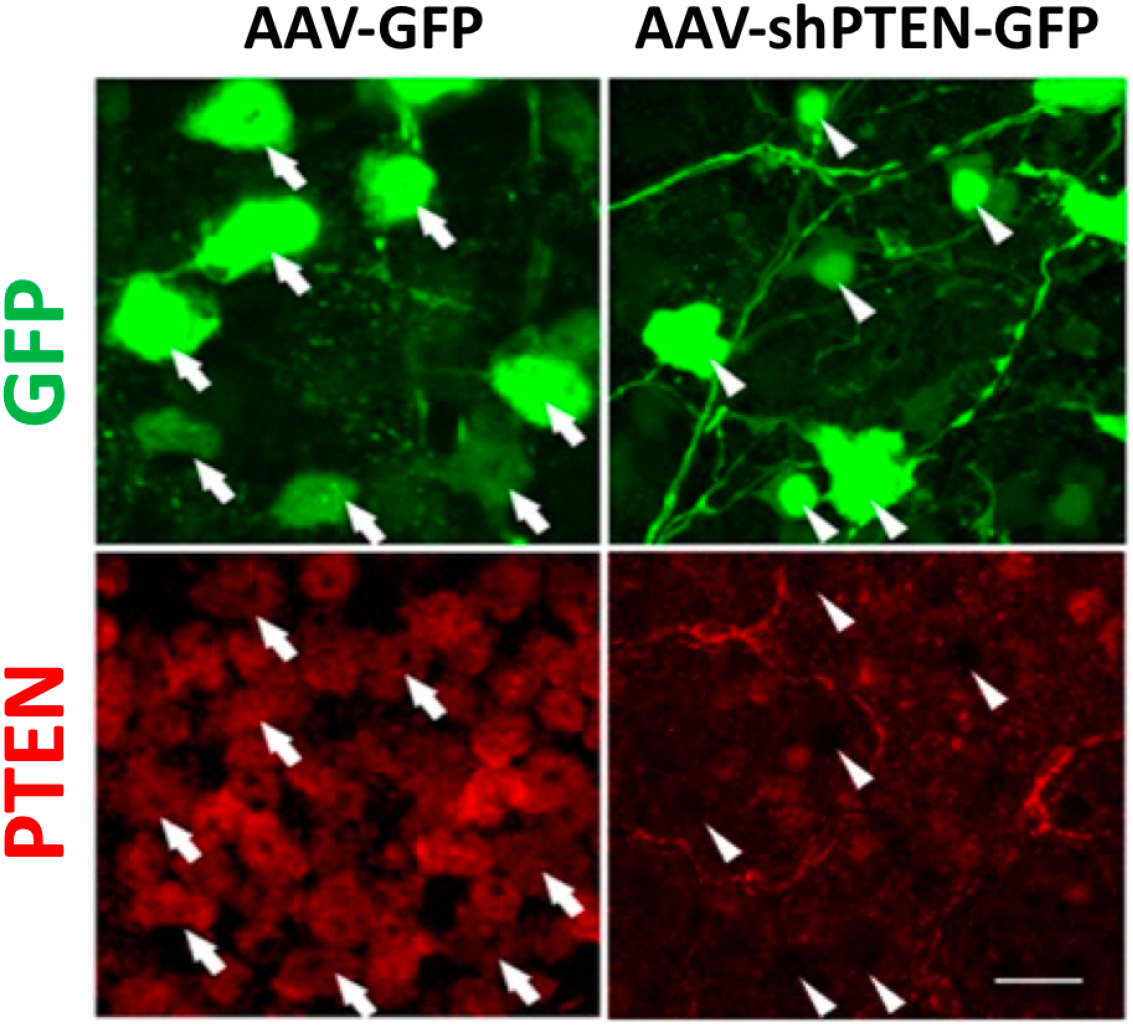
To determine if *Pten* was knocked down by our AAV treatment, we stained the retina for *Pten* (Red) in retinas injected with the AAV-GFP vector (control) or AAV-shPTEN-GFP to knock down *Pten*. In the control retina. all of the GFP positive cells were also well labeled for *Pten*. In the retinas that received the AAV-shPTEN-GFP, the transfected cells (GFP positive) had low levels of *Pten* staining indicating that the vector had in fact decreased the expression of *Pten*.

**Supplementary Table 1.**
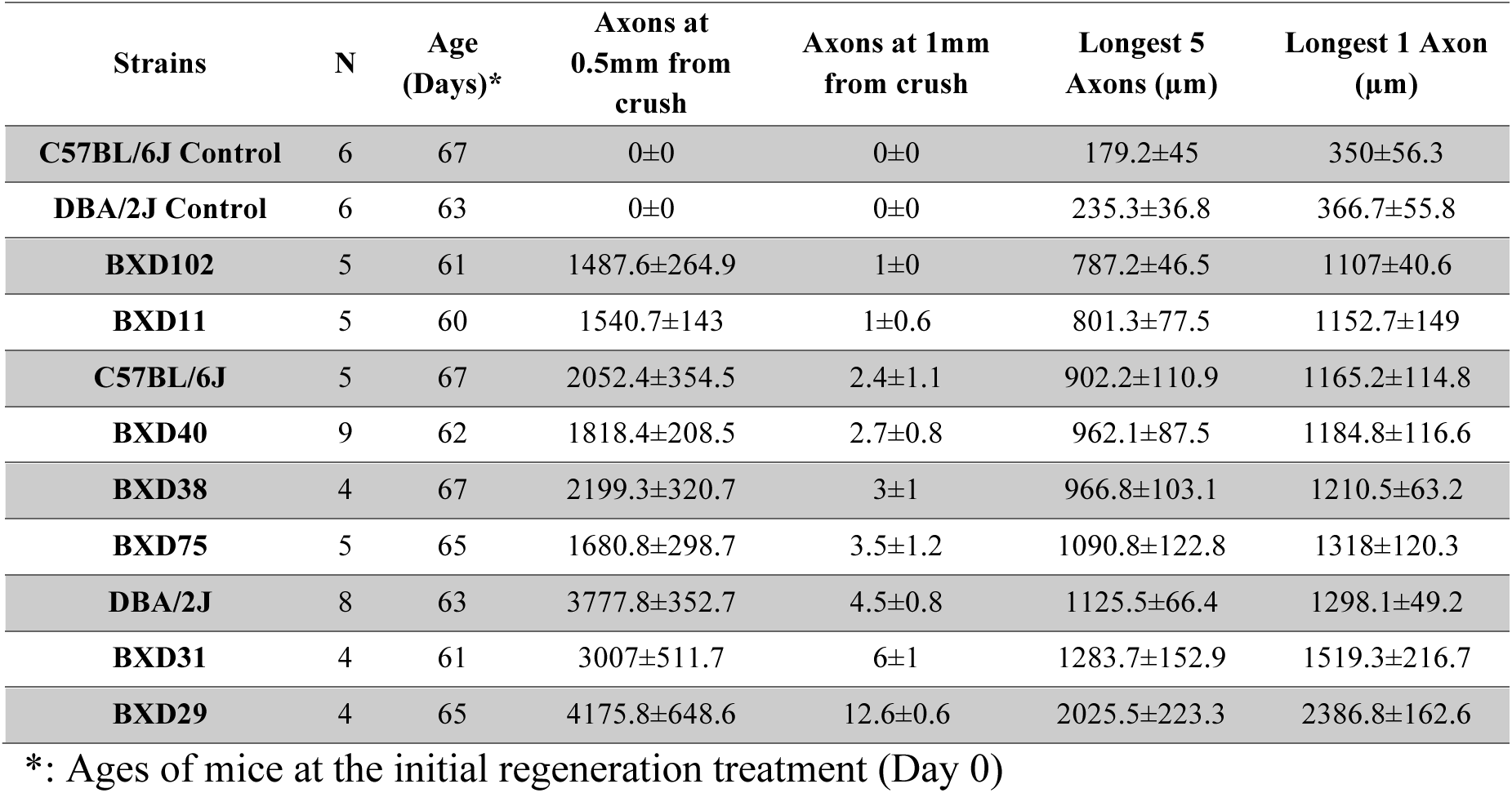
Summary of optic nerve regeneration in the BXD strains

| Strains | N | Age<br>(Days)* | Axons at<br>0.5mm from<br>crush | Axons at 1mm<br>from crush | Longest 5<br>Axons (μm) | Longest 1 Axon<br>(μm) |
| --- | --- | --- | --- | --- | --- | --- |
| **C57BL/6J Control** | 6 | 67 | 0±0 | 0±0 | 179.2±45 | 350±56.3 |
| **DBA/2J Control** | 6 | 63 | 0±0 | 0±0 | 235.3±36.8 | 366.7±55.8 |
| **BXD102** | 5 | 61 | 1487.6±264.9 | 1±0 | 787.2±46.5 | 1107±40.6 |
| **BXD11** | 5 | 60 | 1540.7±143 | 1±0.6 | 801.3±77.5 | 1152.7±149 |
| **C57BL/6J** | 5 | 67 | 2052.4±354.5 | 2.4±1.1 | 902.2±110.9 | 1165.2±114.8 |
| **BXD40** | 9 | 62 | 1818.4±208.5 | 2.7±0.8 | 962.1±87.5 | 1184.8±116.6 |
| **BXD38** | 4 | 67 | 2199.3±320.7 | 3±1 | 966.8±103.1 | 1210.5±63.2 |
| **BXD75** | 5 | 65 | 1680.8±298.7 | 3.5±1.2 | 1090.8±122.8 | 1318±120.3 |
| **DBA/2J** | 8 | 63 | 3777.8±352.7 | 4.5±0.8 | 1125.5±66.4 | 1298.1±49.2 |
| **BXD31** | 4 | 61 | 3007±511.7 | 6±1 | 1283.7±152.9 | 1519.3±216.7 |
| **BXD29** | 4 | 65 | 4175.8±648.6 | 12.6±0.6 | 2025.5±223.3 | 2386.8±162.6 |

\*: Ages of mice at the initial regeneration treatment (Day 0)

